## Supplementary Materials for "Rapid cloning-free mutagenesis of new SARS-CoV-2 variants using a novel reverse genetics platform"

**Figure S1. Clonal virus populations and reconstitution efficiency**

**A** Schematic representation of the workflow to generate clonal virus populations. Right after transfection, cells were diluted to less than 0.5 virus-producing cells/well in 96-well plates. 7 days post-transfection, the supernatant was transferred onto Vero E6-TMPRSS2 cells. Plates were observed until a CPE became apparent.

**B** Clonal virus populations arising from a single virus-producing cell were identified after supernatant transfer onto Vero E6-TMPRSS2 cells. CPE of infectious virus was assessed by microscopy and virus was collected for further analysis. Thereafter, plates were fixed and stained for the fast enumeration of positive wells (in light blue).

**Figure S2. CLEVER primer design for direct mutagenesis**

Shown are the different approaches of primer design in order to guarantee a 100 bp overlap between the neighboring fragments. Primers can either be separated in distance to ensure homology regions or additional nucleotides must be added to the 5' end of the primer. Small nucleotide changes, deletions or insertions can directly be introduced by adjusting the primer annealing sites and the homology regions, as depicted.

**Figure S3. Cloning-free rescue of chimeric virus**

**A** In addition to the chimeric viruses described in Figure 3, the genetic background (region outside of S) was replaced by the sequence of Omicron BA.1 or Omicron BA.5 and combined with a heterologous S sequence of Wuhan, Omicron BA.1 or BA.5, respectively.

**B** Infectious chimeric virus was rescued and assessed via CPE formation on Vero E6 cells. Scale bar represents 100 µm.

**C** The fragments were directly amplified by one-step RT-PCR from viral RNA of Wuhan, Omicron BA.1 and Omicron BA.5. Eight fragments were amplified to prove high flexibility in exchanging fragments.

**D** Scheme for the rapid distinguishment between Wuhan, Omicron BA.1, BA.5 and XBB.1.5 variants. Indicated regions (S or M) were Sanger sequenced to discriminate variants or confirm chimeric viruses. Amino acids are highlighted in yellow for the clear identification of the S gene variant and/or the background (within M).

**Figure S4. Cloning-free rescue of CHIKV and DENV**

**A** Schematic representation of the CHIKV genome and the design for the cloning-free rescue. The genome was divided into three overlapping fragments and a silent SNP was introduced by PCR (red asterisk).

**B and C** Successful recombination of the four PCR products (**B**) within the eukaryotic cell leads to a circular product and virus production (**C**, CPE on BHK-21 cells for rCHIKV and negative control).

**D** Schematic representation of the DENV genome and the design for the cloning-free rescue. The genome has been divided into two overlapping fragments and a silent SNP has been introduced by PCR (red asterisk). The protocol has been tested on two different clinical isolates (DENV1 and DENV3), whereas two different recombination sites were tested for DENV1 (rDENV1-A and rDENV1-B, only one schematically represented).

**E and F** Overlapping PCR products (**E**) were transfected into BHK-21 cells and CPE (**F**) was assessed on VeroE6-TMPRSS2 cells (rDENV1-A, rDENV1-B, rDENV3 and negative control).

Scalebar is 100 $\mu$m in **C** and **F**.

**Table S1. Infectivity assessment of recombinant virus on Vero E6 cells.**

Vero E6 cells were infected with the indicated recombinant virus and pictures were taken 3 days post infection. Of note, the virus was not titrated and the development of CPE is not quantitative. Scale bar represents 100 µm.

**Table S2. Genomic characterization of recombinant SARS-CoV-2 virus based on NGS data.** Mutations with a relative abundance of > 10 % in the entire virus population are listed.

**Table S3. Homology regions successfully used for recombinant SARS-CoV-2, CHIKV and DENV1/DENV3.**

Length, GC content, and the hypothetical annealing temperature (according to OligoCalc, salt-adjusted) are listed. Note that homology regions were chosen independently from GC content or annealing temperature and values are only listed for completion.

**Table S4. Sanger sequencing data of the region of interest for the generated mutant rSARS-CoV-2, rCHIKV and rDENV1/rDENV3.**

**Table S5. Primer tables.**

**Table S6. PCR settings for individual fragments.**

#### Supplementary figure 1: Clonal virus populations and reconstitution efficiency

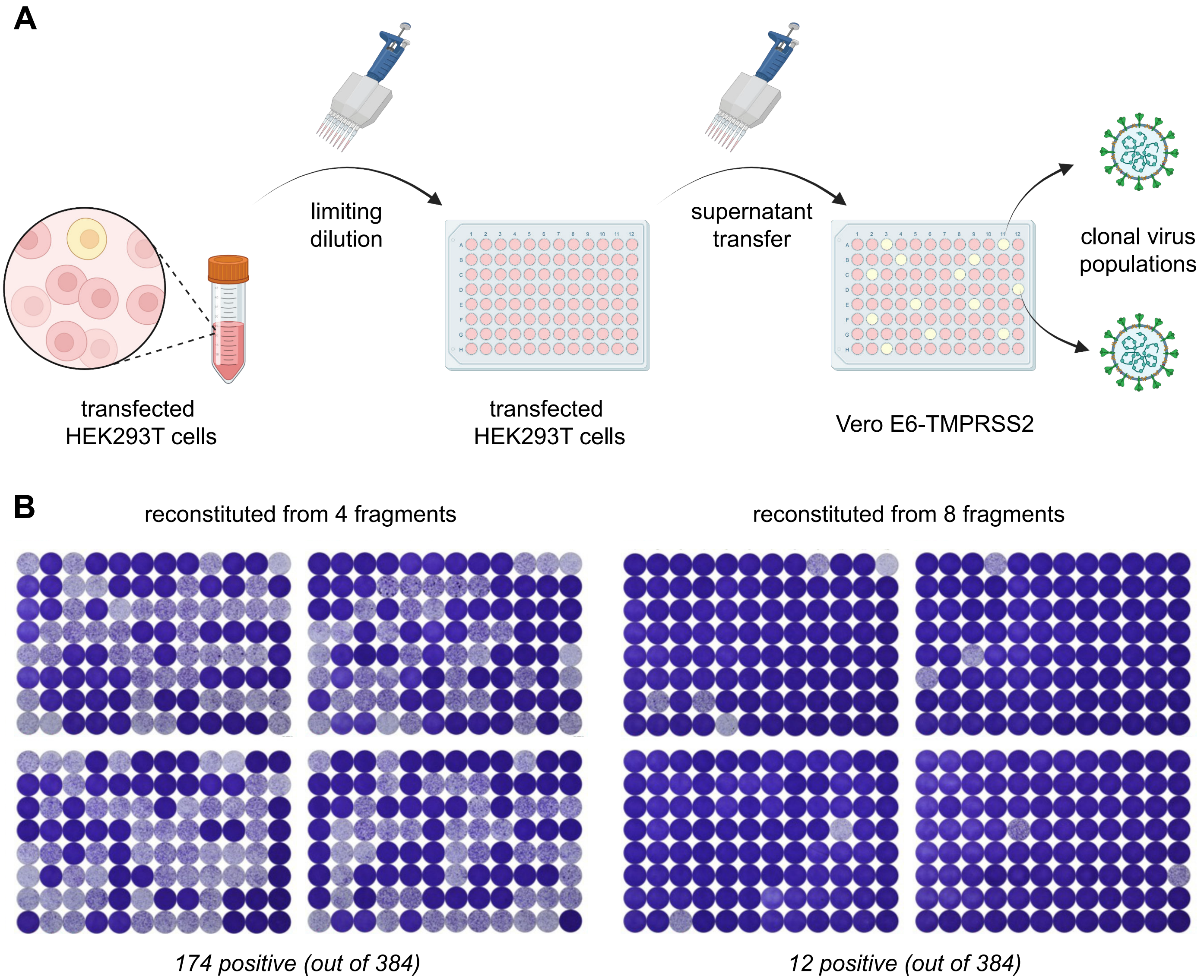

### Supplementary figure 2: CLEVER primer design for direct mutagenesis

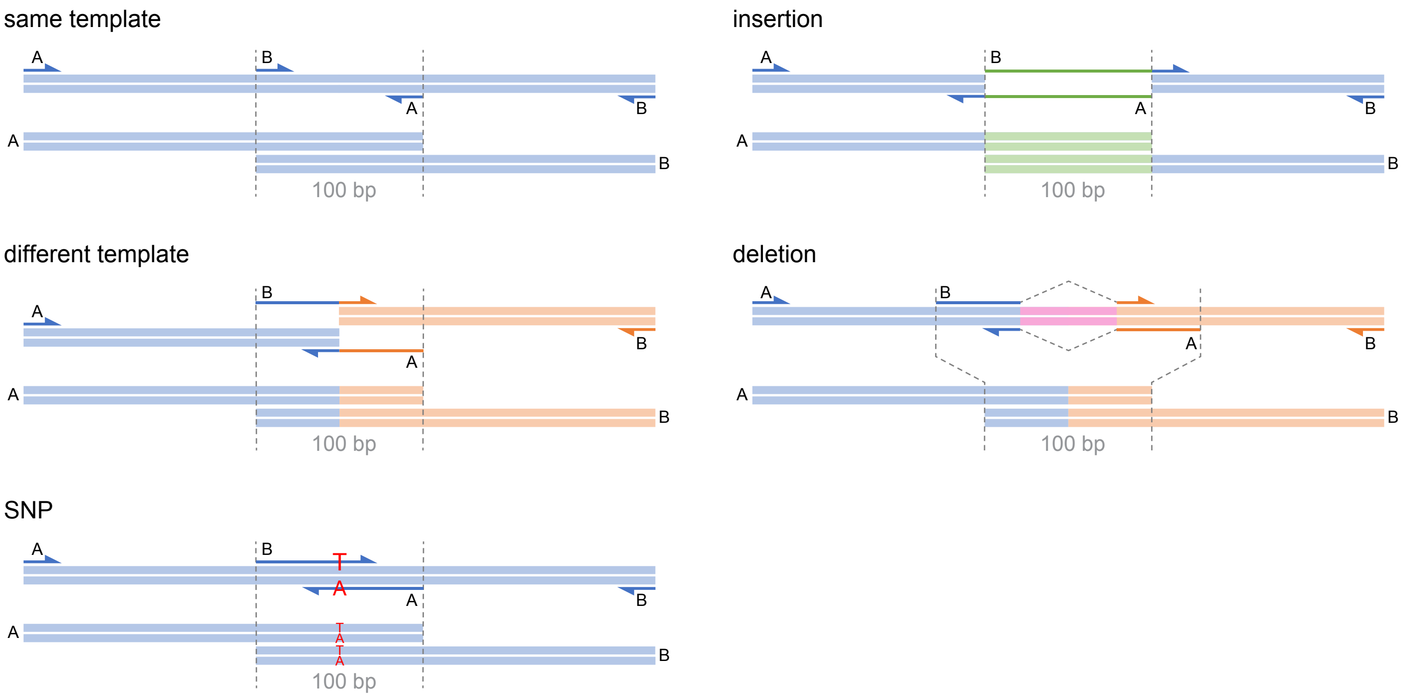

### Supplementary figure 3: Cloning-free rescue of chimeric virus

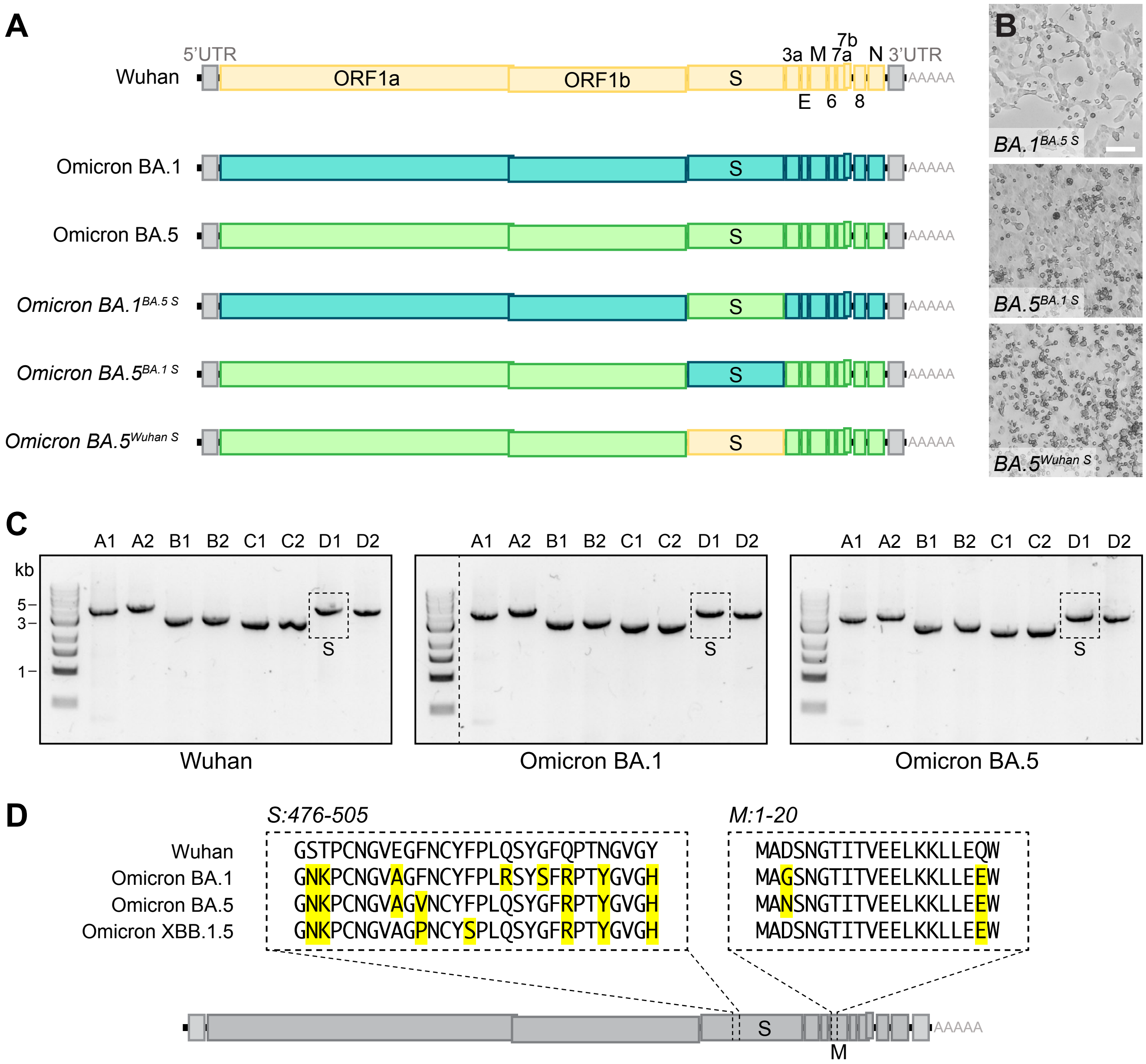

#### Supplementary figure 4: Cloning-free rescue of CHIKV and DENV

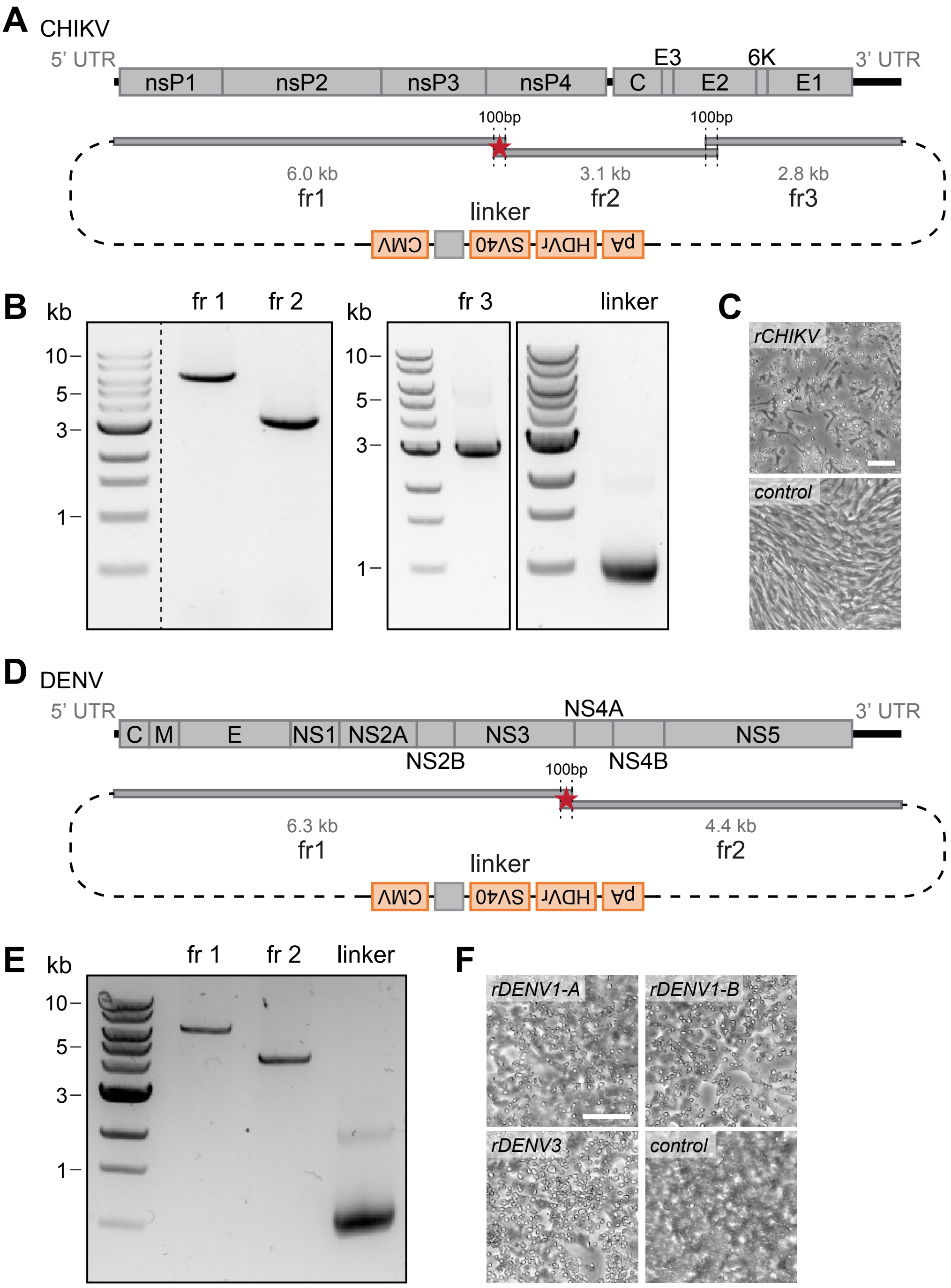

#### Table S1. Infectivity assessment of recombinant virus on Vero E6 cells.

| Number of fragments | Transfection method | Name | Cell line transfected | 3 days post infection on Vero E6 | Description |
|---|---|---|---|---|---|
| - | - | Vero E6 negative control | - | 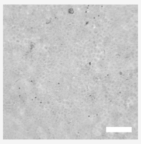 | Uninfected Vero E6 cells. |
| 8 | electroporation | rCOV2-8fr | HEK293T | 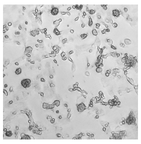 | Rescue from eight fragments. |
| 4 | electroporation | rCOV2-4fr | HEK293T | 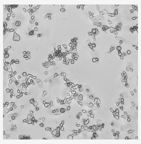 | Rescue from four fragments. |
| 4 | electroporation | rCOV2-293 | HEK293 | 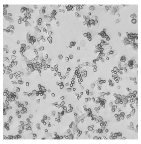 | Rescue from HEK293. |
| 4 | electroporation | rCOV2-CHO | CHO | 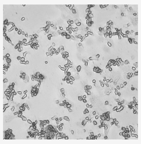 | Rescue from CHO. |
| 4 | electroporation | rCOV2-BHK | BHK-21 | 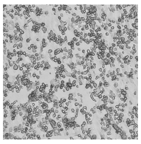 | Rescue from BHK-21. |
| 4 | electroporation | rCOV2-A549 | A549-ACE2 | 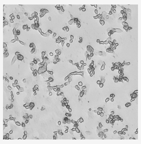 | Rescue from A549-ACE2. |
| 4 | jetPRIME | rCOV2-jP | HEK293T | 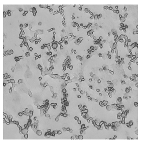 | Rescue after transfection with jetPRIME. |
| 4 | Lipofectamine-3000 | rCOV2-3000 | HEK293T | 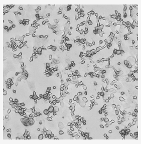 | Rescue after transfection with Lipofectamine-3000. |
| 4 | Lipofectamine-LTX | rCOV2-LTX | HEK293T | 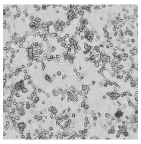 | Rescue after transfection with Lipofectamine-LTX. |

| | | | | | |
|---|---|---|---|---|---|
| 5 | electroporation | rCOV2-500bp | HEK293T | 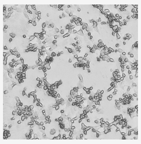 | Rescue including one fragment to be only 500 bp in length. |
| 5 | electroporation | Wuhan$^{BA.1\ S}$ | HEK293T | 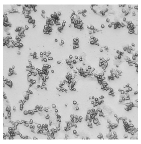 | Chimeric virus, Omicron BA.1 S gene in the background of Wuhan. |
| 5 | electroporation | Wuhan$^{BA.5\ S}$ | HEK293T | 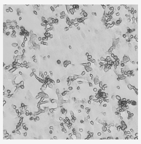 | Chimeric virus, Omicron BA.5 S gene in the background of Wuhan. |
| 5 | electroporation | rCOV2-N501Y | HEK293T | 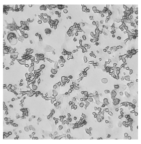 | Direct mutagenesis, incorporation of N501Y substitution in the S gene. |
| 5 | electroporation | rCOV2-G614D | HEK293T | 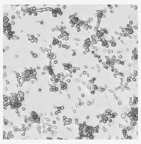 | Direct mutagenesis, incorporation of G614D substitution in the S gene. |
| 5 | electroporation | rCOV2-∆ORF3a | HEK293T | 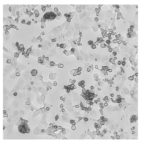 | Direct mutagenesis, deletion of ORF3a. |
| 5 | electroporation | rCOV2-ORF8-FLAG | HEK293T | 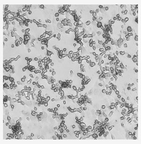 | Direct mutagenesis, incorporation of a 3x-FLAG tag at the C-terminal of ORF8. |
| 9 | electroporation | rCOV2-link-Wuhan | HEK293T | 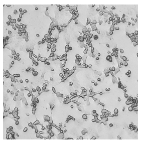 | Rescue from RNA directly using the linker fragment. |
| 9 | electroporation | rCoV2-link-Omicron BA.5$^{BA.1\ S}$ | HEK293T | 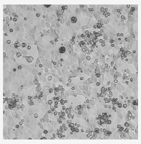 | Rescue of chimeric virus from RNA directly using the linker fragment. Omicron BA.1 S gene in the background of Omicron BA.5. |
| 9 | electroporation | rCoV2-link-Omicron BA.1$^{BA.5\ S}$ | HEK293T | 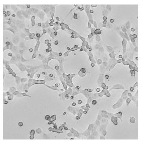 | Rescue of chimeric virus from RNA directly using the linker fragment. Omicron BA.5 S gene in the background of Omicron BA.1. |
| 9 | electroporation | rCoV2-link-Omicron BA.5$^{Wuhan\ S}$ | HEK293T | 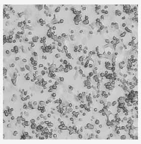 | Rescue of chimeric virus from RNA directly using the linker fragment. Wuhan S gene in the background of Omicron BA.5. |

**Table S2. Genomic characterization of recombinant SARS-CoV-2 virus based on NGS data.** Mutations with a relative abundance of > 10 % in the entire virus population are listed.

| ID | position | | | | | | | "N" reads* | origin of template |
|---|---|---|---|---|---|---|---|---|---|
| | 13551 | 21003* | 21621* | 21638* | 25768* | 28540* | 28967* | | |
| 1 | | K | M | Y | M | | | | T15324C |
| 2 | | | M | Y | M | | | 5979, 9628, 9629, 20231, 20525, 20526 | T15324Y |
| 3 | | | M | Y | M | M | M | | T15324Y |
| 4 | | | M | Y | M | M | | | T15324Y |
| 5 | | | M | Y | M | M | | 5979, 5980, 9628, 9629, 20525, 21002, 21003 | |
| 6 | | K | M | Y | M | M | M | 5979, 5980, 9628, 9629, 20525, 20526 | T15324Y |
| 7 | C to T (13 %) | | M | Y | M | M | | 9628, 9629 | T15324Y |
| 8 | | K | M | Y | M | M | | 5979, 9628, 9629, 20231, 20232, 20526 | T15324Y |
| | *orf1ab* | | *S* | | *orf3a* | *N* | | | |

* Ambiguities or "N" stretches occuring as clusters were not considered as a randomly introduced mutation during the PCR-amplification step but rather as technical artefact or culture adaptation of the virus.

| ID | position | | | | | | | |
|---|---|---|---|---|---|---|---|---|
| | *29*** | *30*** | *79* | *259* | *263* | *264* | 550 | 12464 |
| clonal 1 | | | | | | | | |
| clonal 2 | *A to G* | *A to G* | *A to G* | *A to G* | *A to G* | *A to G* | A to G | |
| clonal 3 | | | | | | | | |
| clonal 4 | | | | | | | | |
| clonal 5 | | | | | | | | G to T |
| | *5'UTR* | | | | | | *orf1ab* | |

*** low coverage (20 reads)*

| ID | position | | | | | | | | | | | | | | |
|---|---|---|---|---|---|---|---|---|---|---|---|---|---|---|---|
| | 3561 | 5834 | 6067 | 18750 | 18897 | 19097 | 20429 | 20763 | 20855 | 20881 | 21137 | 21777 | 25101 | 27787 | 28720 |
| RNA clonal 1 | | T to C | | T to C | | | | | | | A to G | | A to G | T del | C to T |
| RNA clonal 2 | | | | | | | C to T | A to G | G to C | G to A | | | | | |
| RNA clonal 3 | T to C | | T to G | | T to C | C to T | | | | | | G to A | | | |
| | *orf1ab* | | | | | | | | | | | *S* | | *orf7b* | *N* |

**Table S3. Homology regions successfully used for recombinant SARS-CoV-2, CHIKV and DENV1/DENV3**. Length, GC content as well as the hypothetical annealing temperature (according to OligoCalc, salt-adjusted) are listed. Note that homology regions were chosen independently from GC content or annealing temperature and values are only listed for completion.

| | Length (bp) | G/C (%) | Annealing (°C) | Sequence |
|---|---|---|---|---|
| SARS-CoV-2 | 100 | 30 | 83 | GCACTTAAGGGTGGTAAAATTGTTAATAATTGGTTGAAGCAGTTAATTAAAGTTACACTTGTGTTCCTTTTTGTTGCTGCTATTTTCTATTTAATAACAC |
| | 100 | 38 | 86.3 | TATAACTCAAATGAATCTTAAGTATGCCATTAGTGCAAAGAATAGAGCTCGCACCGTAGCTGGTGTCTCTATCTGTAGTACTATGACCAATAGACAGTTT |
| | 100 | 43 | 88.3 | TACAGCTGTTTTAAGACAGTGGTTGCCTACGGGTACGCTGCTTGTCGACTCAGATCTTAATGACTTTGTCTCTGATGCAGATTCAACTTTGATTGGTGAT |
| | 102 | 35 | 85.2 | TGTGTTGAAGAAGTTACAACAACTCTGGAAGAAACTAAGTTCCTCACAGAAAACTTGTTACTTTATATTGACATTAATGGCAATCTTCATCCAGATTCTGCC |
| | 93 | 43 | 87.7 | CACAGGGACTACTCCCACCCAAGAATAGCATAGATGCCTTCAAACTCAACATTAAATTGTTGGGTGTTGGTGGCAAACCTTGTATCAAAGTAG |
| | 107 | 36 | 86 | TTTACAAGTCTTGAAATTCCACGTAGGAATGTGGCAACTTTACAAGCTGAAAATGTAACAGGACTCTTTAAAGATTGTAGTAAGGTAATCACTGGGTTACATCCTAC |
| | 100 | 45 | 89.2 | CAGTCAGCACCTCATGGTGTAGTCTTCTTGCATGTGACTTATGTCCCTGCACAAGAAAAGAACTTCACAACTGCTCCTGCCATTTGTCATGATGGAAAAG |
| | 100 | 46 | 89.6 | GGAACTGTAACTTTGAAGCAAGGTGAAATCAAGGATGCTACTCCTTCAGATTTTGTTCGCGCTACTGCAACGATACCGATACAAGCCTCACTCCCTTTCG |
| | 100 | 34 | 84.6 | CCACGCGAACAAATAGATGGTTATGTCATGCATGCAAATTACATATTTTGGAGGAATACAAATCCAATTCAGTTGTCTTCCTATTCTTTATTTGACATGA |
| | 102 | 37 | 86 | TGCTAGGGAGAGCTGCCTATATGGAAGAGCCCTAATGTGTAAAATTAATTTTAGTAGTGCTATCCCCATGTGATTTTAATAGCTTCTTAGGAGAATGACAAA |
| | 119 | 40 | 88.4 | GTTTAGTGAACCGTATATTAGGTTTATACCTTCCCAGGTAACAAACCAACCAACTTTCGATCTCTTGTAGATCTGTTCTCTAAACGAACTTTAAAATCTGTGTGGCTGTCACTCGGCTG |
| | 100 | 34 | 84.6 | AAGGTTTTAATTGTTACTTTCCTTTACAATCATATGGTTTCCAACCCACTAATGGTGTTGGTTACCAACCATACAGAGTAGTAGTACTTTCTTTTGAACT |
| | 80 | 43 | 86.3 | AACACCAGGAACAAATACTTCTAACCAGGTTGCTGTTCTTTATCAGGGTGTTAACTGCACAGAAGTCCCTGTTGCTATTC |
| | 100 | 43 | 88.3 | ACGACTCTGAGCCAGTGCTCAAAGGAGTCAAATTACATTACACATAAGCACAAGCTGATGAGTACGAACTTATGTACTCATTCGTTTCGGAAGAGACAGG |
| | 84 | 43 | 86.8 | GGCGGAGGTGGTTCTGACTACAAAGACCATGACGGTGATTATAAAGATCATGACATCGATTACAAGGATGACGATGACAAGTAA |
| CHIKV | 100 | 52 | 92 | CCTGTGTACTCGCCTCCGATCAACGTCCGATCGTCCAATCCCGAGTCCGCTGTGGCAGCATGCAATGAGTTCTTAGCTAGAAACTATCCAACTGTCTCAT |
| | 126 | 56 | 95.4 | CGTGCAGAGCACCGCCGCAACTACCGAAGAGATAGAGGTACACATGCCCCCAGACACCCCTGATCGCACATTAATGTCACAACAGTCCGGCAACGTAAAGATCACAGTCAATGGCCAGACGGTGCG |
| | 100 | 31 | 83.4 | AGGGACGTAGGAGATGTTATTTTGTTTTTAATATTTCAAAAAAAAAAAAAAAAAAAAAAAAAAAAAAAAAAAAGGCCGGCATGGTCCCAGCCTCCTCGCT |

| | | | | |
|---|---|---|---|---|
| DENV | 110 | 51 | 92.4 | GGCGTGTACGGTGGGAGGTCTATATAAGCAGAGCTCGTTTAGTGAACCGTATGGCTGCGTGAGACACACGTAGCCTACCAGTTTCTTACTGCTCTACTCTGCAAAGCAAG |
| | 100 | 50 | 91.2 | GGCGTGTACGGTGGGAGGTCTATATAAGCAGAGCTCGTTTAGTGAACCGAGTTGTTAGTCTACGTGGACCGACAAGAACAGTTTCGAATCGGAAGCTTGC |
| | 100 | 59 | 94.9 | TCACCGATCCAGCCAGCATAGCGGCCAGAGGGTACATCTCAACCCGAGTTGGCATGGGTGAAGCAGCTGCGATCTTCATGACAGCCACTCCCCCAGGATC |
| | 100 | 53 | 92.4 | TGACAGCCACTCCCCCAGGATCGGTGGAGGCCTTTCCACAGAGCAATGCCGTTATCCAAGATGAGGAAAGAGACATTCCTGAGAGATCATGGAACTCAGG |
| | 100 | 36 | 85.5 | CCAGAAAATGGAATGGTGCTGTTGAATCAACAGGTTCTAAAAAAAAAAAAAAAAAAAAAAAAAAAAAAAAAAAGGCCGGCATGGTCCCAGCCTCCTCGCT |
| | 100 | 51 | 91.6 | GGCGTGTACGGTGGGAGGTCTATATAAGCAGAGCTCGTTTAGTGAACCGAGTTGTTAGTCTACGTGGACCGACAAGAACAGTTTCGACTCGGAAGCTTGC |
| | 100 | 47 | 90 | CTTGATGCCCGCACTTATTCAGATCCCTTAGCACTCAAGGAATTCAAGGACTTTGCGGCCGGTAGAAAGTCAATCGCCCTTGATCTTGTGACAGAAATAG |
| | 100 | 36 | 85.5 | CCAGAAAATGGAATGGTGCTGTTGAATCAACAGGTTCTAAAAAAAAAAAAAAAAAAAAAAAAAAAAAAAAAAAGGCCGGCATGGTCCCAGCCTCCTCGCT |

#### Table S4. Sanger sequencing data of the region of interest for the generated mutant rSARS-CoV-2, rCHIKV and rDENV1/rDENV3.

| | Mutation/ Region | Sequence |
|---|---|---|
| SARS-CoV-2 | Wuhan(BA.1 S) S | ATGGCAGATTCCAACGGTACTATTACCGTTGAAGAGCTTAAAAAGCTCCTTGAACAATGG |
| | Wuhan(BA.1 S) M | GGTAACAAACCTTGTAATGGTGTTGCAGGTTTTAATTGTTACTTTCCTTTACGATCATATAGTTTCCGACCCACTTATGGTGTTGGTCAC |
| | Wuhan(BA.5 S) S | ATGGCAGATTCCAACGGTACTATTACCGTTGAAGAGCTTAAAAAGCTCCTTGAACAATGG |
| | Wuhan(BA.5 S) M | GGTAACAAACCTTGTAATGGTGTTGCAGGTGTTAATTGTTACTTTCCTTTACGATCATATGGTTTCCAACCCACTTATGGTGTTGGTCAC |
| | rCOV2-N501Y | TACTTTCCTTTACAATCATATGGTTTCCAACCCACTTATGGTGTTGGTTACCAACCATACAGAGTAGTAGTACTTTCTTTTGAACTTCTACATGCACCAGCAACTGTTTGT |
| | rCOV2-N501Y p5 | TACTTTCCTTTACAATCATATGGTTTCCAACCCACTTATGGKGTTGGTTACCACCCATACAGAGTAGTAGTACTTTCTTTTGAACTTCTACATGCACCAGCAACTGTTTGT |
| | rCOV2-G614D | CCAGGAACAAATACTTCTAACCAGGTTGCTGTTCTTTATCAGGATGTTAACTGCACAGAAGTCCCTGTTGCTATTCATGCAGATCAACTTACTCCTACTTGGCGTGTT |
| | rCOV2-G614D p5 | CCAGGAACAAATACTTCTAACCAGGTTGCTGTTCTTTATCAGGATGTTAACTGCACAGAAGTCCCTGTTGCTATTCATGCAGATCAACTTACTCCTACTTGGCGTGTT |
| | rCOV2-ΔORF3a | GACTCTGAGCCAGTGCTCAAAGGAGTCAAATTACATTACACATAAGCACAAGCTGATGAGTACGAACTTATGTACTCATTCGTTTCGGAAGAGACAGGTACGTTAATAGTTAATAGCGTACTTCTTTTT |
| | rCOV2-ORF8-FLAG | ATGAAATTTCTTGTTTTCTTAGGAATCATCACAACTGTAGCTGCATTTCACCAAGAATGTAGTTTACAGTCATGTACTCAACATCAACCATATGTAGTTGATGACCCGTGTCCTATTCACTTCTATTCTAAATGGTATATTAGAGTAGGAGCTAGAAAATCAGCACCTTTAATTGAATTGTGCGTGGATGAGGCTGGTTCTAAATCACCCATTCAGTACATCGATATCGGTAATTATACAGTTTCCTGTTTACCTTTTACAATTAATTGCCAGGAACCTAAATTGGGTAGTCTTGTAGTGCGTTGTTCGTTCTATGAAGACTTTTTAGAGTATCATGACGTTCGTGTTGTTTTAGATTTCATCGGCGGAGGTGGTTCTGACTACAAAGACCATGACGGTGATTATAAAGATCATGACATCGATTACAAGGATGACGATGACAAGTAAACGAACAAACTAAAATGTCT |
| | rCoV2-link-Omicron BA.1(BA.5 S) S | GGTAACAAACCTTGTAATGGTGTTGCAGGTGTTAATTGTTACTTTCCTTTACAATCATATGGTTTCCGACCCACTTATGGTGTTGGTCAC |
| | rCoV2-link-Omicron BA.1(BA.5 S) M | ATGGCAGGTTCCAACGGTACTATTACCGTTGAAGAGCTTAAAAAGCTCCTTGAAGAATGG |
| | rCoV2-link-Omicron BA.5(BA.1 S) S | GGTAACAAACCTTGTAATGGTGTTGCAGGTTTTAATTGTTACTTTCCTTTACGATCATATAGTTTCCGACCCACTTATGGTGTTGGTCAC |
| | rCoV2-link-Omicron BA.5(BA.1 S) M | ATGGCAAATTCCAACGGTAYTATTACCGTTGAAGAGCTTAAAAAGCTCCTTGAAGAATGG |
| | rCoV2-link-Omicron BA.5(Wuhan S) S | GGTAGCACACCTTGTAATGGTGTTGAAGGTTTTAATTGTTACTTTCCTTTACATTCATATGGTTTCCAACCCACTAATGGTGTTGGTTAC |
| | rCoV2-link-Omicron BA.5(Wuhan S) M | ATGGCAAATTCCAACGGTACTATTACCGTTGAAGAGCTTAAAAAGCTCCTTGAAGAATGG |
| | BA.5 Δ3 | ACGACTCTGAGCCAGTGCTCAAAGGAGTCAAATTACATTACACATAAGCACAAGCTGATGAGTACGAACTTATGTACTCATTCGTTTCGGAAGAGATAGGTACGTTAATA |
| | BA.5 Δ3 M | ATGGCAAATTCCAACGGTACTATTACCGTTGAAGAGCTTAAAAAGCTCCTTGAAGAATGG |
| | XBB Δ3 | ACGACTCTGAGCCAGTGCTCAAAGGAGTCAAATTACATTACACATAAGCACAAGCTGATGAGTACGAACTTATGTACTCATTCGTTTCGGAAGAGATAGGTGCGTTAATAAAT |
| | XBB Δ3 M | ATGGCAGATTCCAACGGTACTATTACCGTTGAAGAGCTTAAAAAGCTCCTTGAAGAATGG |
| CHI KV | rCHIKV | GCGCCTGTGTACTCGCCTCCGATCAACGTCCGATTGTCCAATCCCGAGTCCGCTGTGGCAGCATGCAATGAGTTCTTAGCTAGAAACTATCCAACTGTCTCATCATACCAAATTACCGACGAGTATGATGCATATCTAGACATGGTGGACGGGTCGGAGAGTTGCCTGGACCGAGCGACATTCAATCCGTCAAAACTCAGGAGCTACCCGAAACAGCACGCTTACCACGCGCCCTCCATCAGAAGCGCTG |

| | | |
|---|---|---|
| | CHIKV ctr | GCGCCTGTGTACTCGCCTCCGATCAACGTCCGATTGTCCAATCCCGAGTCCGCAGTGGCAGCATGCAATGAGTTCTTAGCTAGAAACTATCCAACTGTCTCATCATACCAAATTACCGACGAGTATGATGCATATCTAGACATGGTGGACGGGTCGGAGAGTTGCCTGGACCGAGCGACATTCAATCCGTCAAAACTCAGGAGCTACCCGAAACAGCACGCTTACCACGCGCCCTCCATCAGAAGCGCTG |
| DENV | rDENV1 A | TACCGATCCAGCTAGCATAGCGGCCAGAGGGTACATCTCAACCCGAGTGGGCATGGGTGAAGCAGCTGCGATCTTTATGACAGCCACTCCCCCAGGATCGGTGGAGGCCTTTCCACAGAGCAATGCCGTTATCCAAGATGAGGAAAGAGACATTCCTGAGAGATCATGGAACTCAGGCTACGACTGGATCACTGACTTTCCAGGTAAAACAGTCTGGTTTGTTCCAAGCATCAAG |
| | DENV1 ctr | TACCGATCCAGCTAGCATAGCGGCCAGAGGGTACATCTCAACCCGAGTGGGCATGGGTGAAGCAGCTGCGATCTTTATGACAGCCACTCCCCCAGGATCGGTGGAGGCCTTTCCACAGAGCAATGCAGTCATCCAAGATGAGGAAAGAGACATTCCTGAGAGATCATGGAACTCAGGCTACGACTGGATCACTGACTTTCCAGGTAAAACAGTCTGGTTTGTTCCAAGCATCAAG |
| | rDENV1 B | TATGGATGAAGCACATTTTACCGATCCAGCCAGCATAGCGGCCAGAGGGTACATCTCAACCCGAGTTGGCATGGGTGAAGCAGCTGCGATCTTTATGACAGCCACTCCCCCAGGATCGGTGGAGGCCTTTCCACAGAGCAATGCAGTCATCCAAGATGAGGAAAGAGACATTCCTGAGAGATCATGGAACTCAGGCTACGACTGGATCACTGACTTTCCAGGTAAAACAGTCTGGTTTGTTCCAAGCATCAAG |
| | rDENV3 | GTAGCATCAGAAGGGATCAAATATACAGATAGGAAATGGTGCTTTGATGGACAGCGCAACAACCAAATTTTAGAGGAAAACATGGACGTGGAAATCTGGACAAAGGAAGGAGAAAAGAAAAAATTAAGACCTAGGTGGCTTGATGCCCGCACTTATTCAGATCCCTTAGCACTCAAGGAATTCAAGGACTTTGCGGCCGGTAGAAAGTCAATCGCCCTTGATCTTGTGACAGAAATAGGAAGAGTGCCTTCACACCTAGCCCACAGAACGAGAAACGCTCTGGACAATT |
| | DENV3 ctr | GTAGCATCAGAAGGGATCAAATATACAGATAGGAAATGGTGCTTTGATGGACAGCGCAACAACCAAATTTTAGAGGAAAACATGGACGTGGAAATCTGGACAAAGGAAGGAGAAAAGAAAAAATTAAGACCTAGGTGGCTTGATGCCCGCACTTATTCAGATCCCTTAGCACTCAAGGAATTCAAGGACTTTGCGGCTGGTAGAAAGTCAATCGCCCTTGATCTTGTGACAGAAATAGGAAGAGTGCCTTCACACCTAGCCCACAGAACGAGAAACGCTCTGGACAATT |

### Table S5. Primer tables.

| | Fragment | Primer | Sequence 5’ to 3’ | Amplicon Size |
|---|---|---|---|---|
| SARS-CoV-2 | A1 | F CMV | CGATGTACGGGCCAGATATACG | 4694 |
| | | R A1-A2 | GGCAGAATCTGGATGAAGATTGCC | |
| | A2 | F A2-A1 | TGTGTTGAAGAAGTTACAACAACTCTG | 4672 |
| | | R A-B | GTGTTATTAAATAGAAAATAGCAGCAACAAAAAGGAACACAAGTGT AACTTTAATTAACTGCTTCAACC | |
| | B1 | F B-A | GCACTTAAGGGTGGTAAAATTGTTAATAATTGGTTGAAGCAGTTAAT TAAAGTTACACTTGTGTTCC | 3292 |
| | | R B1-B2 | CTACTTTGATACAAGGTTTGCCACC | |
| | B2 | F B2-B1 | CACAGGGACTACTCCCACCC | 3415 |
| | | R B-C | AAACTGTCTATTGGTCATAGTACTACAGATAGAGACACCAGCTACG GTGCGAGCTCTATTCTTTGCAC | |
| | C1 | F C-B | TATAACTCAAATGAATCTTAAGTATGCCATTAGTGCAAAGAATAGAG CTCGCACCGTAGCTGGTG | 3048 |
| | | R C1-C2 | GTAGGATGTAACCCAGTGATTACC | |
| | C2 | F C2-C1 | TTTACAAGTCTTGAAATTCCACGTAGG | 3006 |
| | | R C-D | ATCACCAATCAAAGTTGAATCTGCATCAGAGACAAAGTCATTAAGAT CTGAGTCGACAAGCAGCG | |
| | D1 | F D-C | TACAGCTGTTTTAAGACAGTGGTTGCCTACGGGTACGCTGCTTGTC GACTCAGATCTTAATGACTTTGTC | 4622 |
| | | R D1-D2 | CGAAAGGGAGTGAGGCTTGTATC | |
| | D2 | F D2-D1 | GGAACTGTAACTTTGAAGCAAGGTG | 4707 |
| | | R SV40 | GCGGCCGCCAGACATGATAAG | |
| | A | F CMV | CGATGTACGGGCCAGATATACG | 9264 |
| | | R A-B | GTGTTATTAAATAGAAAATAGCAGCAACAAAAAGGAACACAAGTGT AACTTTAATTAACTGCTTCAACC | |
| | B | F B-A | GCACTTAAGGGTGGTAAAATTGTTAATAATTGGTTGAAGCAGTTAAT TAAAGTTACACTTGTGTTCC | 6614 |
| | | R B-C | AAACTGTCTATTGGTCATAGTACTACAGATAGAGACACCAGCTACG GTGCGAGCTCTATTCTTTGCAC | |
| | C | F C-B | TATAACTCAAATGAATCTTAAGTATGCCATTAGTGCAAAGAATAGAG CTCGCACCGTAGCTGGTG | 5947 |
| | | R C-D | ATCACCAATCAAAGTTGAATCTGCATCAGAGACAAAGTCATTAAGAT CTGAGTCGACAAGCAGCG | |
| | D | F D-C | TACAGCTGTTTTAAGACAGTGGTTGCCTACGGGTACGCTGCTTGTC GACTCAGATCTTAATGACTTTGTC | 9229 |
| | | R SV40 | GCGGCCGCCAGACATGATAAG | |
| | D1 614 | F D-C | | 2535 |

| | | | | |
|---|---|---|---|---|
| | | R G614D | GAATAGCAACAGGGACTTCTGTGCAGTTAACATCCTGATAAAGAAC AGCAACCTGGTTAGAAGTATTTGTTCC | |
| | D2 614 | F G614D | AACACCAGGAACAAATACTTCTAACCAGGTTGCTGTTCTTTATCAG GATGTTAACTGCACAGAAGTCCCTGTTGCTATTC | 6774 |
| | | R SV40 | | |
| | D1 501 | F D-C | | 2212 |
| | | R N501Y | AGTTCAAAAGAAAGTACTACTACTCTGTATGGTTGGTAACCAACAC CATAAGTGGGTTGGAAACCATATG | |
| | D2 501 | F N501Y | AAGGTTTTAATTGTTACTTTCCTTTACAATCATATGGTTTCCAACCCA CTTATGGTGTTGGTTACCAACCATAC | 7117 |
| | | R SV40 | | |
| | D1 Δorf3a | F D-C | | 4495 |
| | | R Δorf3a | CCTGTCTCTTCCGAAACGAATGAGTACATAAGTTCGTACTCATCAG CTTGTGCTTATGTGTAATGTAATTTGACTCC | |
| | D2 Δorf3a | F Δorf3a | ACGACTCTGAGCCAGTGCTCAAAGGAGTCAAATTACATTACACATA AGCACAAGCTGATGAGTACGAACTTATGTACTC | 3912 |
| | | R SV40 | | |
| | D1 orf8-3xFLAG | F D-C | | 7356 |
| | | R orf8-3xFLAG | TTACTTGTCATCGTCATCCTTGTAATCGATGTCATGATCTTTATAATC ACCGTCATGGTCTTTGTAGTCAGAACCACCTCCGCCGATGAAATCT AAAACAACACG | |
| | D2 orf8-3xFLAG | F orf8-3xFLAG | GGCGGAGGTGGTTCTGACTACAAAGACCATGACGGTGATTATAAA GATCATGACATCGATTACAAGGATGACGATGACAAGTAAACGAACA AACTAAAATGTCTG | 1873 |
| | | R SV40 | | |
| | linker | F linker | AGGCCACGCGGAGTACGATCGAGTGTACAGTGAACAATGCTAGGG AGAGCTGCC<br>or<br>TGCTAGGGAGAGCTGCC (equal performance) | 1106 |
| | | R linker | CAGCCGAGTGACAGCCACAC | |
| | A1-linker | F A-linker | GTTTAGTGAACCGTATATTAGGTTTATACCTTCCCAGGTAACAAACC | 4083 |
| | | R A1-A2 | | |
| | D2-linker | F D2-D1 | | 4451 |
| | | R D-linker | TTTGTCATTCTCCTAAGAAGCTATTAAAATCACATG | |
| | D1 500 | F D-C | | 500 |
| | | R 500 | TCATGTCAAATAAAGAATAGGAAGACAACTG | |
| | D2 500 | F 500 | CCACGCGAACAAATAGATGGTTATG | 8829 |
| | | R SV40 | | |

| | | | | |
|---|---|---|---|---|
| CHIKV | fr1 | F fr1-linker CHIKV | GGCGTGTACGGTGGGAGGTCTATATAAGCAGAGCTCGTTTAGTGAA CCGTATGGCTGCGTGAGACACACG | 6128 |
| | | R fr1-fr2 CHIKV | ATG AGA CAG TTG GAT AGT TTC TAG CTA AGA ACT CAT TGC ATG CTG CCA CAG CGG ACT CGG GAT TGG AC | |
| | fr2 | F fr2_fr1 CHIKV | CCT GTG TAC TCG CCT CCG ATC AAC GTC CGA TCG TCC AAT CCC GAG TCC GCT GTG GCA GCA TGC AAT GAG | 3157 |
| | | R fr2-fr3 CHIKV | CGCACCGTCTGGCCATTGACTGTGATCTTTACGTTGCCGGACTGTT GTGACATTAATGTGCGATCAGGGGTG | |
| | fr3 | F fr3-fr2 CHIKV | CGTGCAGAGCACCGCCGCAACTACCGAAGAGATAGAGGTACACAT GCCCCCAGACACCCCTGATCGCAC | 2816 |
| | | R fr3-linker CHIKV | AGCGAGGAGGCTGGGACCATGCCGGCCTTTTTTTTTTTTTTTTTTTT TTTTTTTTTTTTTTTTGAAATATTAAAAACAAAATAACATCTCC | |
| | linker CHIKV | F linker-fr3 CHIKV | AGGGACGTAGGAGATGTTATTTTGTTTTTAATATTTCAAAAAAAAAAA AAAAAAAAAAAAAAAAAAAAAAAAAGGCCGGCATGGTCCCAG | 1009 |
| | | R linker-fr1 CHIKV | CTTGCTTTGCAGAGTAGAGCAGTAAGAAACTGGTAGGCTACGTGTG TCTCACGCAGCCATACGGTTCACTAAACGAGC | |

| | | | | |
|---|---|---|---|---|
| DENV | linker DENV1 | F linker-fr2_DENV1 | CCA GAA AAT GGA ATG GTG CTG TTG AAT CAA CAG GTT CTA AAA AAA AAA AAA AAA AAA AAA AAA AAA AAA AAA AGG CCG GCA TGG TCC CAG | 989 |
| | | R linker-fr1_DENV1 | GCA AGC TTC CGA TTC GAA ACT GTT CTT GTC GGT CCA CGT AGA CTA ACA ACT CGG TTC ACT AAA CGA GC | |
| | fr1 mA | F fr1-linker_DENV1 | GGC GTG TAC GGT GGG AGG TCT ATA TAA GCA GAG CTC GTT TAG TGA ACC GAG TTG TTA GTC TAC GTG GAC C | 5612 |
| | | R fr1-fr2 DENV1 A | CCT GAG TTC CAT GAT CTC TCA GGA ATG TCT CTT TCC TCA TCT TGG ATA ACG GCA TTG CTC TGT GGA AAG GCC TCC AC | |
| | fr2 mA | F fr2-fr1 DENV1 A | TGA CAG CCA CTC CCC CAG GAT CGG TGG AGG CCT TTC CAC AGA GCA ATG CCG TTA TCC AAG ATG AGG AAA GAG | 5335 |
| | | R fr2-linker_DENV1 | AGCGAGGAGGCTGGGACCATGCCGGCCTTTTTTTTTTTTTTTTTTTT TTTTTTTTTTTTTTTTAGAACCTGTTGATTCAACAGCACCATTCCATTTT CTGGCGTTCTGTGCCTGGAATGAT | |
| | fr1 mB | F fr1-linker_DENV1 | GGC GTG TAC GGT GGG AGG TCT ATA TAA GCA GAG CTC GTT TAG TGA ACC GAG TTG TTA GTC TAC GTG GAC C | 5534 |
| | | R fr1-fr2 DENV1 B | GAT CCT GGG GGA GTG GCT GTC ATG AAG ATC GCA GCT GCT TCA CCC ATG CCA ACT CGG GTT GAG ATG TAC CCT C | |
| | fr2 mB | F fr2-fr1 DENV1 B | TCA CCG ATC CAG CCA GCA TAG CGG CCA GAG GGT ACA TCT CAA CCC GAG TTG GCA TGG GTG AAG CAG CTG C | 5413 |
| | | R fr2-linker_DENV1 | AGCGAGGAGGCTGGGACCATGCCGGCCTTTTTTTTTTTTTTTTTTTT TTTTTTTTTTTTTTTTagaacctgttgattcaacagcaccattccattttctggcgttctgtgcctg gaatgat | |
| | linker DENV3 | F linker-fr2 DENV3 | CCA GAA AAT GGA ATG GTG CTG TTG AAT CAA CAG GTT CTA AAA AAA AAA AAA AAA AAA AAA AAA AAA AAA AAA AGG CCG GCA TGG TCC CAG | 989 |
| | | R linker-fr1 DENV3 | GCA AGC TTC CGA GTC GAA ACT GTT CTT GTC GGT CCA CGT AGA CTA ACA ACT CGG TTC ACT AAA CGA GC | |
| | fr1 | F fr1-linker DENV3 | GGC GTG TAC GGT GGG AGG TCT ATA TAA GCA GAG CTC GTT TAG TGA ACC GAG TTG TTA GTC TAC GTG GAC C | 6450 |

| | | | | |
|---|---|---|---|---|
| | | R fr1-fr2 DENV3 | CTA TTT CTG TCA CAA GAT CAA GGG CGA TTG ACT TTC TAC CGG CCG CAA AGT CCT TGA ATT CCT TGA GTG CTA AGG GAT CTG AAT AAG | |
| | fr2 | F fr2-fr1 DENV3 | CTT GAT GCC CGC ACT TAT TCA GAT CCC TTA GCA CTC AAG GAA TTC AAG GAC TTT GCG GCC GGT AGA AAG TCA ATC GCC CTT GAT CTT GTG AC | 4468 |
| | | R fr2-linker_DENV3 | AGCGAGGAGGCTGGGACCATGCCGGCCTTTTTTTTTTTTTTTTTTTTTTTTTTTTTTTTTTTTTTagaacctgttgattcaacagcaccattccattttctggcgttctgtgcctggaatgat | |

#### Table S6. PCR settings for individual fragments.

| | Fragment | Cycling Conditions | Elongation |
|---|---|---|---|
| SARS-CoV-2 | A | 5 cycles at 57 °C | 4 min 40 s |
| | | 20 cycles at 65 °C | |
| | B | 5 cycles at 47 °C | 3 min 20 s |
| | | 20 cycles at 65 °C | |
| | C | 5 cycles at 54 °C | 3 min |
| | | 20 cycles at 65 °C | |
| | D | 5 cycles at 50 °C | 4 min 20 s |
| | | 20 cycles at 65 °C | |
| | A1 | 5 cycles at 52 °C | 2 min 30 s |
| | | 20 cycles at 64 °C | |
| | A2 | 5 cycles at 52 °C | 2 min 30 s |
| | | 20 cycles at 64 °C | |
| | B1 | 5 cycles at 48 °C | 1 min 45 s |
| | | 20 cycles at 63 °C | |
| | B2 | 5 cycles at 48 °C | 1 min 45 s |
| | | 20 cycles at 63 °C | |
| | C1 | 5 cycles at 48 °C | 1 min 45 s |
| | | 20 cycles at 63 °C | |
| | C2 | 25 cycles at 61 °C | 1 min 45 s |
| | D1 | 25 cycles at 63 °C | 2 min 30 s |
| | D2 | 5 cycles at 52 °C | 2 min 30 s |
| | | 20 cycles at 64 °C | |
| | linker | 25 cycles at 58 °C | 35 s |
| CHIKV | linker CHIKV | 5 cycles at 54 °C | 30 s |
| | | 20 cycles at 64 °C | |
| | fr1 CHIKV | 25 cycles at 64 °C | 3 min 10 s |
| | fr2 CHIKV | 25 cycles at 62 °C | 1 min 40 s |
| | fr3 CHIKV | 25 cycles at 61 °C | 1 min 30 s |

| | | | |
|---|---|---|---|
| DENV | linker DENV1 | 5 cycles at 52 °C | 30 s |
| | | 20 cycles at 62 °C | |
| | fr1 DENV1 A | 5 cycles at 55 °C | 2 min 50 s |
| | | 20 cycles at 62 °C | |
| | fr2 DENV1 A | 5 cycles at 50 °C | 2 min 50 s |
| | | 20 cycles at 62 °C | |
| | fr1 DENV1 B | 5 cycles at 55 °C | 2 min 50 s |
| | | 20 cycles at 62 °C | |
| | fr1 DENV1 B | 5 cycles at 50 °C | 2 min 50 s |
| | | 20 cycles at 62 °C | |
| | linker DENV3 | 5 cycles at 52 °C | 30 s |
| | | 20 cycles at 62 °C | |
| | fr1 DENV3 | 5 cycles at 50 °C | 3 min 20 s |
| | | 20 cycles at 62 °C | |
| | fr2 DENV3 | 5 cycles at 50 °C | 2 min 20 s |
| | | 20 cycles at 62 °C | |
